# A PDZ-RapGEF promotes synaptic development in *C. elegans* through a Rap/Rac signaling pathway

**DOI:** 10.1101/2025.05.27.656318

**Authors:** Reagan Lamb, Michael Scales, Julie Watkins, Martin Werner, Salvatore J Cherra

## Abstract

Small G proteins coordinate the development of nerve terminals. The activity of G proteins is finely tuned by GTPase regulatory proteins. Previously, we observed that PXF-1, a *Caenorhabditis elegans* GTPase regulatory protein, is required for the function of cholinergic motor neurons. Here, we investigated how PXF-1 coordinates the development of presynaptic terminals at the molecular level. We observed that PXF-1 acts through RAP-1 to promote synapse development. Subsequently, we found that *pxf-1* mutants display a reduction in RAC-2 activity, which is required for cholinergic synapse development. We observed that RAC-2 acts downstream of RAP-1. Finally, we identified a physical interaction between RAP-1 and TIAM-1, a Rac guanine exchange factor, which links PXF-1 function to the presynaptic actin cytoskeleton through RAC-2 activation. These findings highlight how small G protein signaling pathways interact to coordinate the development of presynaptic terminals.

## Introduction

Synapses mediate the communication between presynaptic neurons and their postsynaptic targets, such as other neurons or muscle cells. During development, presynaptic terminals accumulate the necessary equipment for neurotransmission, like active zone scaffolds, voltage-gated calcium channels, synaptic vesicles, and exocytosis machinery. The formation of active zones and clustering of synaptic vesicle pools require the assembly of presynaptic actin filaments (Chia et al., 2014; Chia et al., 2012; Zhang and Benson, 2001). One set of proteins that modulate actin filaments is the family of GTPases known as small guanine nucleotide-binding proteins (small G proteins). The family of small GTPases is comprised of five subfamilies: Rho, Ras, Arf, Rab, and Ran. Some subfamily members, like Rho, Rac, and Rap, modulate the actin cytoskeleton. In mammalian neurons, Rac proteins influence dendrite development, synapse function, and memory (Cheng et al., 2021; Rao et al., 2019). In the *C. elegans* nervous system, Rac homologs regulate axon guidance and promote synaptic vesicle clustering (Lundquist et al., 2001; Stavoe and Colón-Ramos, 2012). Other members, like Rap proteins, ensure proper neuronal migration, synaptic transmission, and plasticity (Franco et al., 2011; Jossin and Cooper, 2011; Morozov et al., 2003; Pan et al., 2008; Subramanian et al., 2013). In *C. elegans,* the Rap homolog RAP-2 restricts synapse formation to ensure proper tiling of presynaptic sites in motor neurons (Chen et al., 2018a). Overall, small G proteins play essential roles during nervous system development.

The activity of small G proteins is coordinated by GTPase activating proteins (GAPs) and guanine nucleotide exchange factors (GEFs). In general, GAPs reduce G protein signaling by stimulating the hydrolysis of GTP by the G protein, which leads to its inactivation. GEFs accelerate the exchange of GDP for GTP, which stimulates G protein signaling. RasGAPs can act on both Ras and Rap to limit their signaling activity and are essential for synapse formation, dendrite development, and synaptic plasticity (Araki et al., 2020; Chen et al., 2018a; Duan et al., 2014; Vazquez et al., 2004). PDZ-containing RapGEFs promote Rap signaling to ensure proper neuronal development and synaptic function. For example, RapGEF2 or RapGEF6 knockout mice display deficits in neural progenitor development (Maeta et al., 2016). Separately, RapGEF6 was shown to modulate neuronal activity and synaptic plasticity in various brain regions (Levy et al., 2015). In *C. elegans*, we found that the PDZ-GEF, PXF-1, is required for synaptic development through an actin-mediated mechanism (Lamb et al., 2022).

In this study, we further investigated how PXF-1 functions to modulate the abundance of synaptic vesicles at presynaptic terminals using the *C. elegans* neuromuscular junction (NMJ). We found that mutations in two small G proteins, *rap-1* or *rac-2*, reduced synapse development in cholinergic motor neurons, like *pxf-1* mutants. Activating mutations in either RAP-1 or RAC-2 were sufficient to restore synapse development in *pxf-1* mutants. Additional data indicate that RAP-1 acts upstream of RAC-2 and its GEF, TIAM-1, to promote synapse development. This study has uncovered a PXF-1 signaling pathway that sequentially activates two small G proteins to maintain presynaptic actin filaments and promote the accumulation of synaptic vesicles at cholinergic terminals.

## Results

### PXF-1 functions in cholinergic neurons to promote the accumulation of synaptic vesicles during development

We previously observed that PXF-1 is required for normal levels of synaptic vesicle markers in cholinergic neurons at the neuromuscular junction (Lamb et al., 2022). Since *pxf-1* mutants produce synapses that contain a dimmer fluorescent signal from synaptic vesicle markers, we sought to determine if this reduction in synaptic vesicle markers was associated with a delay in synapse development. Using mCherry::RAB-3 expressed in cholinergic neurons, we observed increased fluorescence during development in wild type and *pxf-1* animals (**Figure 1A-B**). However, the mCherry::RAB-3 signal was much lower in *pxf-1* mutants as compared to wild type animals (**Figure 1A-B**). We observed a lower fluorescence signal as early as L2 animals that reached statistical significance by L3 and L4 stages.

**Figure 1.**
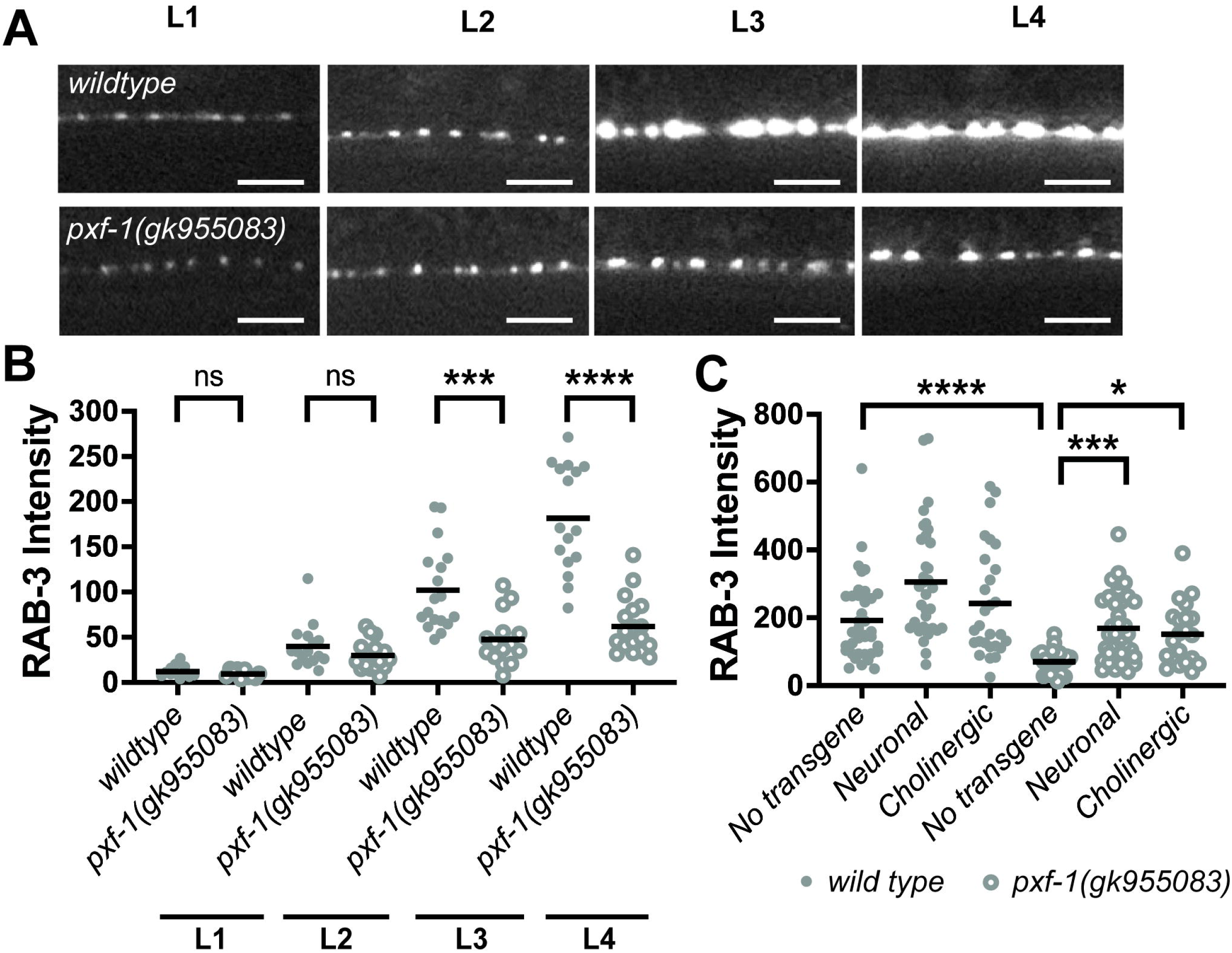
Mutant *pxf-1* animals display a deficiency in the accumulation of RAB-3 labeled vesicles during development. **(A)** Representative images of mCherry::RAB-3 labeled synapses in the dorsal cord of wildtype and *pxf-1(gk955083)* mutant animals during larval stages: L1, L2, L3, and L4. Scale bars indicate 5 μm. **(B)** Quantification of mCherry::RAB-3 grayscale fluorescence intensity. Gray dots represent individual animals, black bars indicate means. n = 12-18. ***p<0.001; ****p<0.0001; ns = not significant. **(C)** Quantification of mCherry::RAB-3 grayscale fluorescence intensity in wild type and *pxf-1(gk955083)* mutants expressing *pxf-1* cDNA in all neurons, only cholinergic neurons, or no transgene. For pan-neuronal expression, *bluEx53* and *bluEx54* analyses were combined into a single dataset named Neuronal, and for cholinergic expression, *bluEx59* and *bluEx61* analyses were combined into a single dataset named Cholinergic. Filled gray dots represent individual wild type animals, open dots represent *pxf-1(gk955083)* mutant animals, and black bars indicate means. n = 21-37. *p<0.05; ***p<0.001; ****p<0.0001; ns = not significant.

We previously observed that PXF-1 functions in neurons but not muscle to promote neuromuscular function (Lamb et al., 2022). To determine in which neurons PXF-1 functions to promote the accumulation of mCherry::RAB-3-labeled synaptic vesicles, we first expressed wild type *pxf-1* cDNA in all neurons using the *rgef-1* promoter. We found that pan-neuronal expression of *pxf-1* cDNA was sufficient to increase mCherry::RAB-3 in *pxf-1* mutant animals (**Figure 1C**). To determine if *pxf-1* expression solely in cholinergic neurons is sufficient to restore proper synapse development, we used the *unc-17b* promoter to drive *pxf-1* expression. We observed that expression of *pxf-1* in cholinergic neurons also was sufficient to increase cholinergic RAB-3::mCherry levels in *pxf-1* mutant animals (**Figure 1C**). Together, these data suggest that PXF-1 functions in a cell autonomous manner to promote the development of neuromuscular junctions.

### RAP-1 activation restores synaptic vesicle intensity in *pxf-1* mutants

To determine which G proteins mediate the effects of PXF-1 on synapse development, we next investigated the Rap homologs *rap-1* and *rap-2*, which function with PXF-1 to promote epithelial development (Pellis-van Berkel et al., 2005). We measured the intensity and density of cholinergic NMJs labeled by mCherry::RAB-3 in *rap-1* and *rap-2* mutant animals. We found that only *rap-1* mutants displayed a decrease in mCherry::RAB-3 intensity, but *rap-2* mutants were indistinguishable from wild type animals **(Figure 2A-D)**. We also observed a small but significant increase in RAB-3-labeled puncta in *rap-1* mutant animals **(Figure 2E)**.

**Figure 2.**
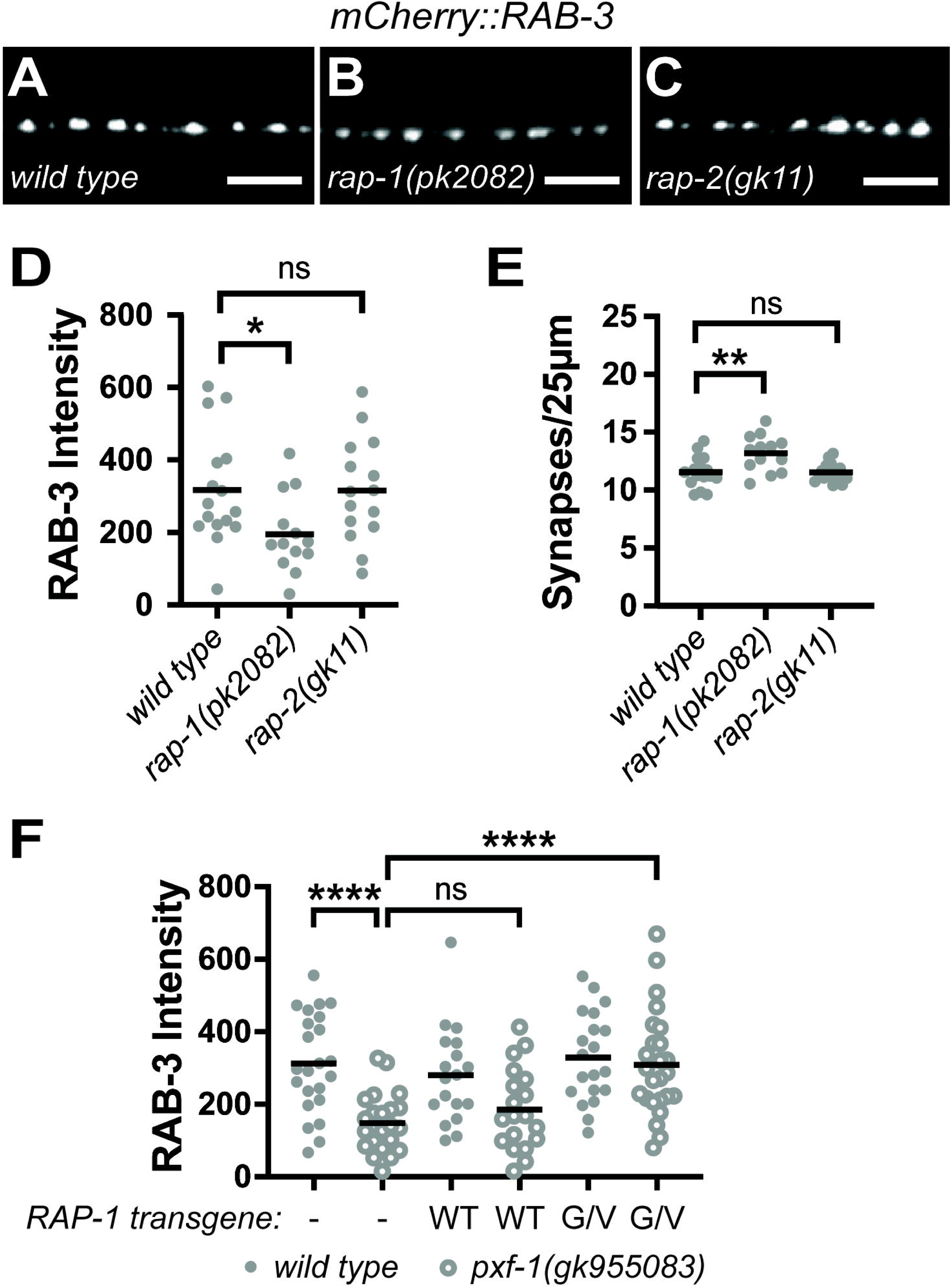
RAP-1 activation restores synaptic vesicle intensity in *pxf-1* mutants. **(A-C)** Representative images of mCherry::RAB-3 labeled cholinergic synapses in the dorsal cord of **(A)** *wild type,* **(B***) rap-1(pk2082),* **(C)** *rap-2(gk11).* Scale bars are 5 μm. **(D)** Quantification of mCherry::RAB-3 grayscale fluorescence intensity. **(E)** Quantification of synaptic density. Gray dots represent individual animals, black bars indicate means. n = 13-16. *p<0.05; **p<0.01; ns = not significant. **(F)** Quantification of mCherry::RAB-3 grayscale fluorescence intensity from wild type or *pxf-1(gk955083)* mutant animals expressing no transgene labeled as a dash, RAP-1(WT)::mGFP labeled as WT, or RAP-1(G12V)::mGFP labeled as G/V transgenes in cholinergic motor neurons. For RAP-1(WT), *bluEx143* and *bluEx144* were combined into one dataset, and for RAP-1(G12V), *bluEx146* and *bluEx147* were combined into one dataset. Each data point represents an individual animal. Filled dots are wild type animals and open dots are *pxf-1(gk955083)* mutants. Black bars indicate means. n = 18-25. ****p<0.0001; ns = not significant.

If RAP-1 is the target of PXF-1, we would expect that RAP-1 signaling would be reduced in *pxf-*1 mutants. Therefore, we hypothesized that activating RAP-1 signaling in cholinergic neurons should restore mCherry::RAB-3 intensity in *pxf-1* mutant animals. To increase RAP-1 signaling in cholinergic neurons, we expressed a constitutively active RAP-1(G12V) cDNA under the *unc-17b* promoter in the *pxf-1(gk955083)* mutants expressing mCherry::RAB-3. This constitutively active mutation blocks hydrolysis of the bound GTP and prevents the G protein from returning to an inactive state (Hammond et al., 2015). To control for increased levels of expression of RAP-1 in the transgenic animals, we also generated a cholinergic RAP-1(WT) cDNA transgene. We found that expression of the constitutively active RAP-1(G12V) cDNA was able to restore the intensity of mCherry::RAB-3 in *pxf-1* mutant animals to wild type levels **(Figure 2F**). However, expression of the RAP-1(WT) cDNA transgene did not increase the intensity of RAB-3::mCherry-labeled puncta in *pxf-1* mutant animals (**Figure 2F**). These data indicate that exogenous activation of RAP-1 signaling can bypass the requirement for PXF-1 for synapse development, suggesting that RAP-1 functions downstream of PXF-1.

### Mutations in *rac-2* reduce aldicarb sensitivity, synaptic vesicle intensity, and presynaptic F-actin levels

We previously found that neuronal expression of WVE-1 was sufficient to restore synapse development in *pxf-1* mutants (Lamb et al., 2022); however, WVE-1 is not a known effector molecule for RAP-1. Rather, the Rac subfamily of Rho G proteins are known to act through the WAVE/SCAR complex to regulate actin filament formation (**Figure 3A**) (Westphal et al., 2000). To determine which Rac homologs may be involved in PXF-1 signaling, we measured the aldicarb sensitivity of Rac mutants. We found that two mutations, *ced-10(n1993)* and *rac-2(ok326),* caused a significant decrease in aldicarb sensitivity (**Figure 3B**). Next, we examined synaptic vesicle abundance in *ced-10* and rac-2 mutant animals by measuring endogenous UNC-17 levels using immunofluorescence. We found that *ced-10(n1993), rac-2(ok326),* and *rac-2(gk281)* mutant animals exhibited decreases in UNC-17 intensity, suggesting a decrease in synaptic vesicle abundance (**Figure 3C-G**).

**Figure 3.**
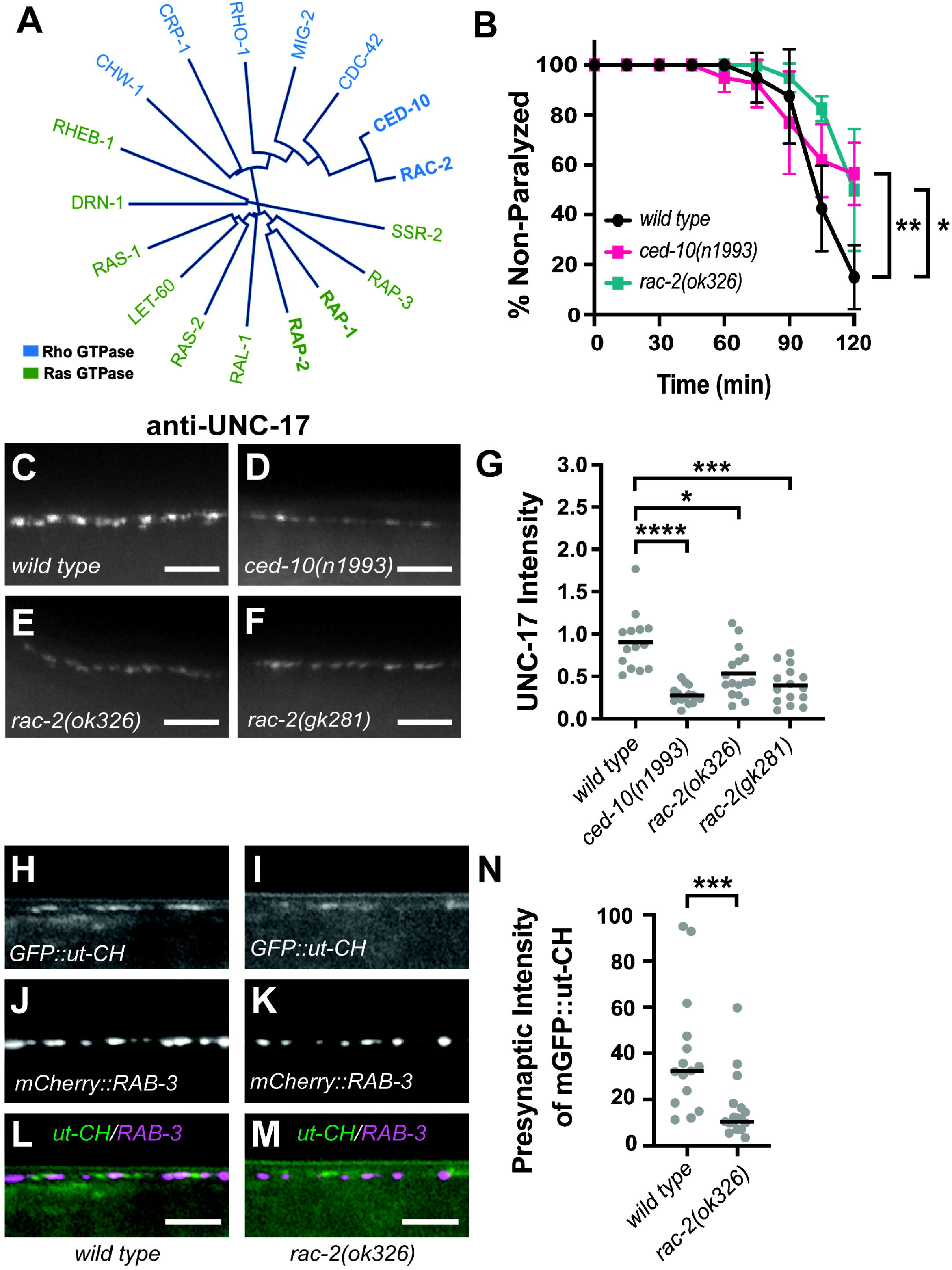
Mutations in *rac-2* reduce aldicarb sensitivity, synaptic vesicle intensity, and presynaptic F-actin levels. **(A)** Phylogenetic tree of Rho and Ras G protein family members. **(B)** Quantification of non-paralyzed animals during exposure to 1 mM aldicarb. Symbols represent means; error bars represent SEM. n = 5 trials of 10 animals each. *p<0.05; **p<0.01. **(C-F)** Representative images of dorsal cord stained by UNC-17 antibodies in **(C)** *wild type,* **(D)** *ced-10(n1993),* **(E)** *rac-2(ok326),* and **(F)** *rac-2(gk281)*. Scale bars are 5 μm. **(G)** Quantification of fluorescence intensity of UNC-17 staining in dorsal nerve cord. Gray dots represent individual animals, black bars indicate means. n = 14-16. *p<0.05; ***p<0.001; ****p<0.0001. **(H-M)** Representative images of dorsal cord in wild type and *rac-2(ok326)* mutant animals with **(H,I)** mGFP labeled calponin homology domain of F-actin binding protein utrophin (GFP::ut-CH), **(J,K)** mCherry labeled RAB-3 in cholinergic synapses, and **(L,M)** merged channels. Scale bars are 5 μm. **(N)** Quantification of grayscale intensity of mGFP::ut-CH within 1 μm of the peak of mCherry::RAB-3 fluorescence signal. Gray dots represent individual animals, black bars indicate median. n= 15-17, ***p<0.001.

To understand how PXF-1 interacts with Rac GTPase signaling, we chose to focus on RAC-2 for the remainder of this study. To determine whether *rac-2* modulates the actin cytoskeleton, like *pxf-1* (Lamb et al., 2022), we measured presynaptic F-actin in cholinergic neurons using mGFP::ut-CH to label filamentous actin and the mCherry::RAB-3 marker to visualize cholinergic synapses. We observed a decrease in mGFP::ut-CH signal in mCherry::RAB-3-labeled synaptic terminals in *rac-2* mutant animals (**Figure 3H-N**), suggesting that RAC-2 promotes the assembly or maintenance of presynaptic actin filaments. Overall, these findings indicate that RAC-2 acts like PXF-1 to promote synapse development in cholinergic neurons in an actin-based manner.

### RAC-2 and PXF-1 act in the same pathway

While it is evident that *rac-*2 mutant animals phenocopy *pxf-1* mutants, it is still unclear whether they are working together to coordinate synapse development. To investigate whether RAC-2 acted in the same pathway as PXF-1, we used animals expressing mCherry::RAB-3 in cholinergic neurons. We found that animals containing mutations in *pxf-1* or *rac-2* displayed a decrease in synaptic vesicle intensity compared to wild type animals but no change in synapse density (**Figure 4A-F**). Additionally, *pxf-1(gk955083); rac-2(ok326)* double mutants showed a decrease in mCherry::RAB-3 intensity that was not significantly different from either single mutant (**Figure 4A-E**), indicating that PXF-1 and RAC-2 act in the same pathway.

**Figure 4.**
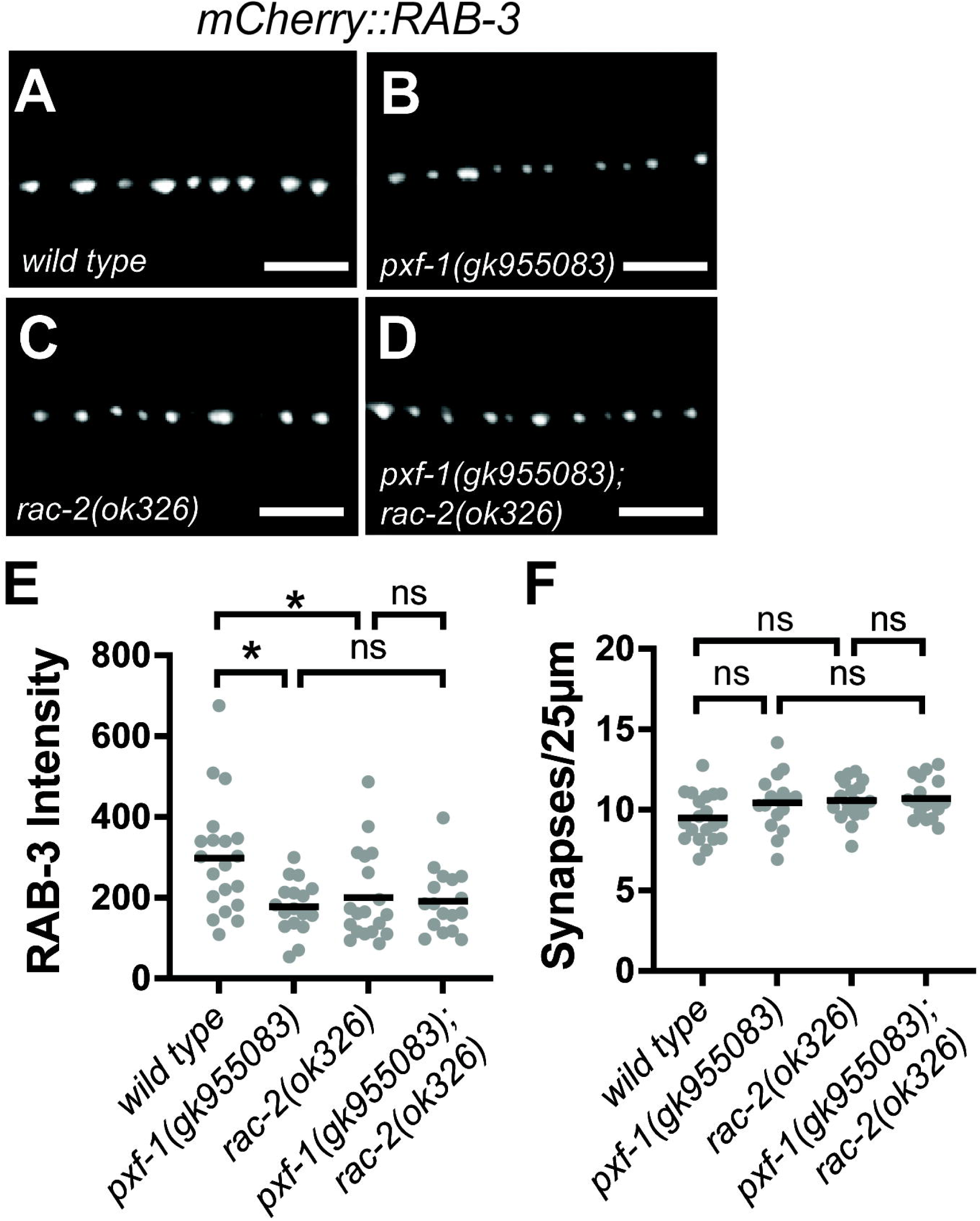
RAC-2 and PXF-1 act in the same pathway. **(A-D)** Representative images of mCherry::RAB-3 labeled cholinergic synapses in the dorsal cord of **(A)** wild type, **(B)** *pxf-1(gk955083),* **(C)** *rac-2(ok326)* and **(D)** *pxf-1(gk955083); rac-2(ok326)* animals. Scale bars are 5 μm. **(E)** Quantification of mCherry::RAB-3 grayscale fluorescence intensity. Gray dots represent individual animals, black bars indicate means. n = 17-20. *p<0.05; ns = not significant. (F) Quantification of synapse density. Gray dots represent individual animals, black bars indicate means. n = 17-20. ns = not significant.

### PXF-1 modulates RAC-2 activity

To determine if RAC-2 activity levels in cholinergic motor neurons were influenced by PXF-1 function, we expressed a fluorescence resonance energy transfer (FRET)-fluorescence lifetime imaging microscopy (FLIM) biosensor under the *unc-17b* promoter. Based on previous designs (Aoki and Matsuda, 2009; Chen et al., 2018b; Graham et al., 2001), our biosensor expresses mCherry fused to the Rac binding domain (RBD) from *C. elegans pak-1,* a T2A ribosomal skip sequence, and mGFP fused to *rac-2* cDNA (**Figure 5A**). When RAC-2 is activated, mGFP::RAC-2 binds to mCherry::RBD, which produces FRET between the mGFP donor and the mCherry acceptor and reduces the time mGFP exists in an excited state.

**Figure 5.**
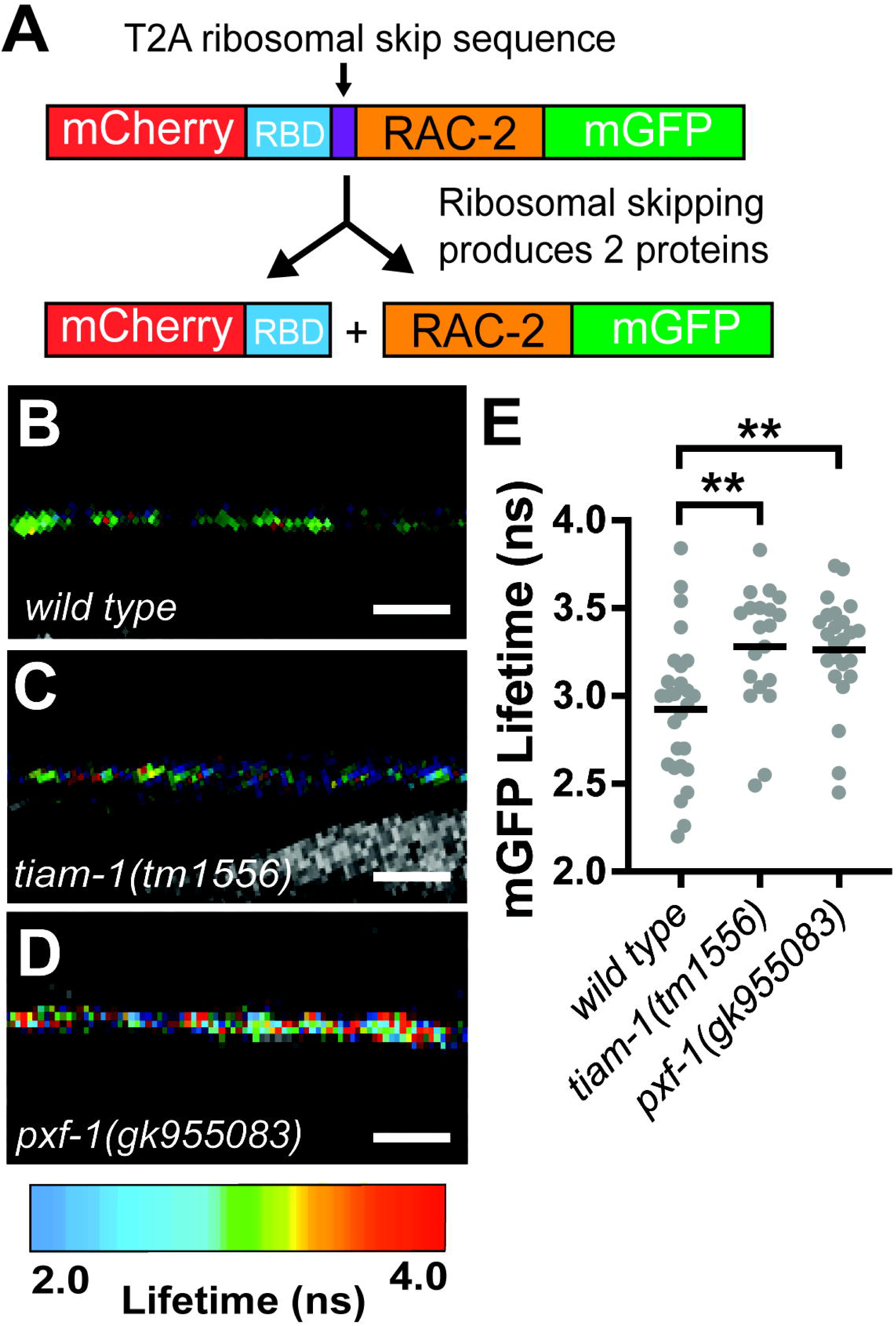
PXF-1 modulates RAC-2 activity. **(A)** Schematic of biosensor plasmid containing mCherry::RBD and RAC-2::mGFP. **(B-D)** FastFLIM images of mGFP in the dorsal cord of **(B)** wild type, **(C)** *tiam-1(tm1556)*, and **(D)** *pxf-1(gk955083)* animals expressing the biosensor in cholinergic neurons. mGFP lifetime has been pseudo-colored to represent fluorescence lifetime values in nanoseconds. Scale bar is 5 μm. **(E)** Mean lifetime fluorescence of RAC-2::mGFP in wild type*, tiam-1(tm1556)* and *pxf-1(gk955083)* mutant backgrounds. Independent alleles for the RAC-2 biosensor (*bluEx71* and *bluEx72*) were used for each genotype; graphs show data that were pooled from *bluEx71* and *bluEx72*. Gray dots represent individual animals, black bars indicate means. n = 18-25. **p<0.01.

We first tested whether our sensor was able to detect changes in RAC-2 activity. In neurons, Rac proteins are activated through their canonical Rac GEF, TIAM-1 (Brar et al., 2022; Demarco et al., 2012); therefore, we compared the lifetime of mGFP::RAC-2 between wild type and *tiam-1(tm1556)* mutant animals. We found that the mean lifetime fluorescence of mGFP::RAC-2 in *tiam-1(tm1556)* animals was higher than wild type animals, indicating a decrease in RAC-2 activity, which is consistent with TIAM-1 function (**Figure 5B-E**). Additionally, these changes in lifetime fluorescence are within the range of changes caused by biologically relevant events (Stubbs et al., 2005).

To determine if *pxf-1* mutants affected the level of RAC-2 activity, compared mGFP::RAC-2 lifetime fluorescence to wild type animals. Similar to *tiam-1* mutant animals, we observed a higher mean lifetime for mGFP::RAC-2 in *pxf-1* mutants as compared to wild type animals (**Figure 5B-E**). This indicates PXF-1 promotes the activation of RAC-2 signaling in cholinergic neurons, which is decreased in *pxf-1* mutant animals, and suggests that RAC-2 may function downstream of PXF-1.

### RAC-2 acts downstream of PXF-1

To determine if RAC-2 acts downstream of PXF-1, we examined whether activation of RAC-2 was sufficient to rescue synaptic vesicle defects. We expressed RAC-2(WT) or the constitutively activated RAC-2(G12V) cDNA in cholinergic neurons and measured the intensity of mCherry::RAB-3 in the wild type and *pxf-1(gk955083)* mutant animals. Like RAP-1(G12V) mutant transgenes, expression of RAC-2(G12V) was sufficient to increase synaptic vesicle intensity in *pxf-1* mutants to wild type levels (**Figure 6A**). Unexpectedly, exogenous expression of RAC-2(WT) also increased mCherry::RAB-3 intensity in *pxf-1* mutants. Together, these data suggest that RAC-2 signaling functions downstream of PXF-1 to promote synapse development; however, our data with the RAC-2(WT) transgene suggest that additional factors may regulate RAC-2 activity.

**Figure 6.**
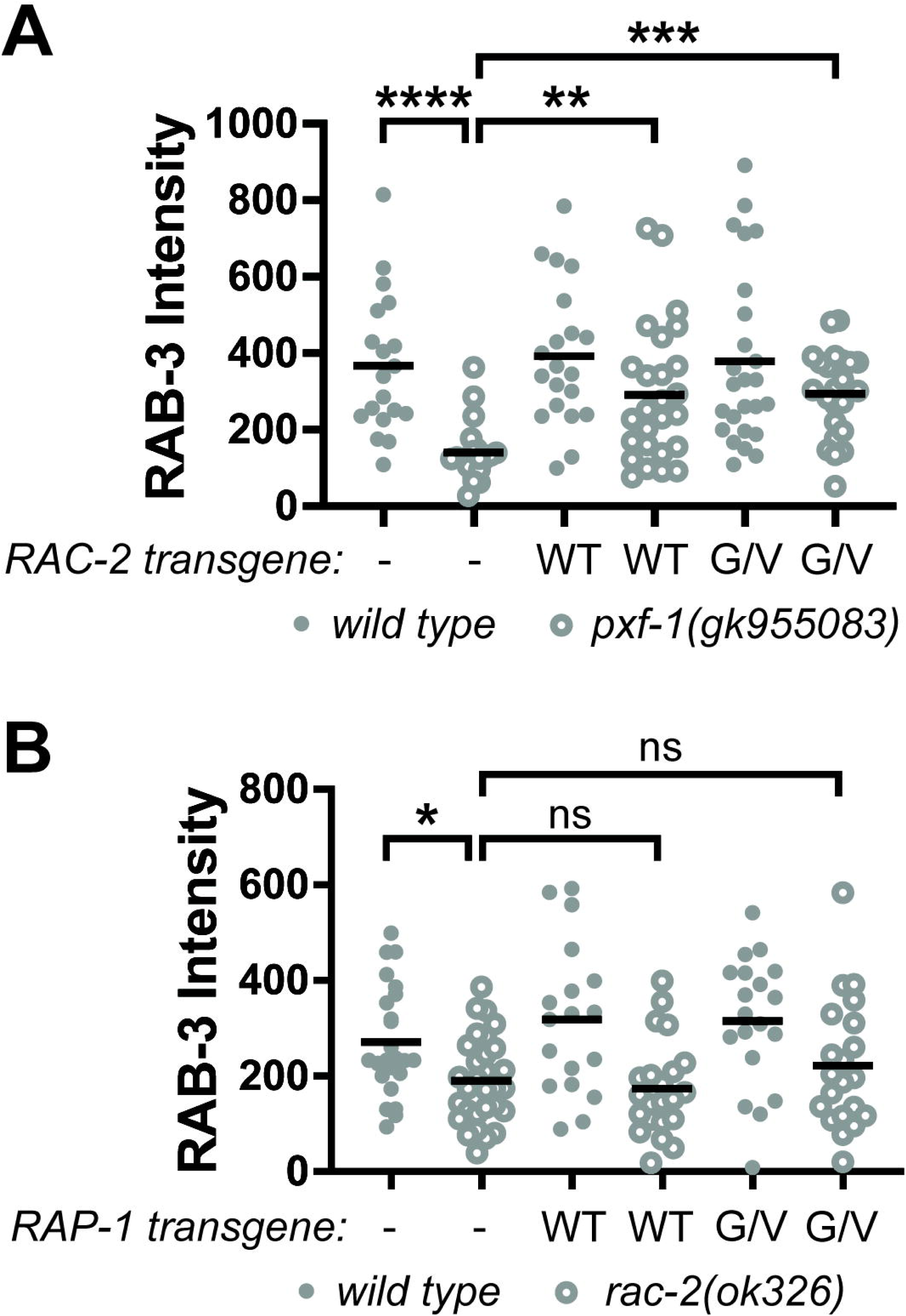
RAC-2 functions downstream of PXF-1 and RAP-1 to promote synapse development. **(A)** Quantification of mCherry::RAB-3 grayscale fluorescence intensity from wild type or *pxf-1(gk955083)* mutant animals expressing no transgene labeled as a dash, RAC-2(WT)::mGFP labeled as WT, or RAC-2(G12V)::mGFP labeled as G/V transgenes. For RAC-2(WT), *bluEx166* and *bluEx168* were combined into one dataset, and for RAC-2(G12V), *bluEx169* and *bluEx171* were combined into one dataset. Each data point represents an individual animal. Filled gray dots are wild type and open dots are *pxf-1(gk955083)* mutant animals. Black bars indicate means. n = 19-26. **p<0.01; ***p<0.001. **(B)** Quantification of mCherry::RAB-3 grayscale fluorescence intensity from wild type or *pxf-1(gk955083)* mutant animals expressing no transgene labeled as a dash, RAP-1(WT)::mGFP labeled as WT, or RAP-1(G12V)::mGFP labeled as G/V transgenes. For RAP-1(WT), *bluEx143* was used, and for RAP-1(G12V), *bluEx146* was used. Each data point represents an individual animal. Filled gray dots are wild type and open dots are *rac-2(ok326)* mutant animals. Black bars indicate means. n = 18-30. *p<0.05; ns = not significant.

### RAP-1 functions upstream of RAC-2 to promote synapse development

Activation of RAP-1 or RAC-2 was sufficient to restore synapse development in *pxf-1* mutant animals (**Figures 2F and 6A)**. While these data indicate that RAP-1 and RAC-2 function in the same pathway as PXF-1, they do not provide insight into the hierarchy of these two G proteins in the signaling pathway. To determine the order of how RAP-1 and RAC-2 function to modulate synaptic development, we measured mCherry::RAB-3 levels in *rac-2* mutants expressing the RAP-1(WT) or RAP-1(G12V) transgene in cholinergic motor neurons. Expression of RAP-1(WT) or RAP-1(G12V) was not able to restore mCherry::RAB-3 levels in *rac-2* mutants to wild type levels (**Figure 6B**). Since the same RAP-1(G12V) transgenes were sufficient to restore synapse development in *pxf-1* mutants but did not rescue synapse development in *rac-2* mutants, these data indicate that RAP-1 functions downstream of PXF-1 but upstream of RAC-2.

### TIAM-1 functions with PXF-1 to restore synapse development

We next sought to determine how PXF-1 and RAP-1 signaling influences RAC-2 activity. TIAM1 and VAV3 were identified as Rac GEFs that modulate synapse development or function (Duman et al., 2013; Ulc et al., 2017; Um et al., 2014). The *C. elegans* genome has homologs for both TIAM1, called *tiam-1*, and VAV3, called *vav-1* (Demarco et al., 2012; Duman et al., 2013). Since *vav-1* mutants cause embryonic lethality, hypersensitivity to aldicarb, and function in an interneuron to control locomotor behavior (Etheridge et al., 2015; Fry et al., 2014), we decided to focus on *tiam-1*, which is required for RAC-2 activation (**Figure 5E**). We measured mCherry::RAB-3 intensity in cholinergic neurons in two *tiam-1* single mutants (*tm1556* or *ok772*) and in double mutants with *pxf-1*. We found that both *tiam-1* alleles decreased mCherry::RAB-3 intensity (**Figure 7A-E and 7G-K)**. Double mutants between *tiam-1* and *pxf-1* displayed a reduction in mCherry::RAB-3 intensity, but there were no significant differences between the single and double mutants (**Figure 7A-E and 7G-K**). There were no changes in synapse number in any single or double mutants **(Figure 7F and 7L)**, indicating that TIAM-1 and PXF-1 act in the same pathway.

**Figure 7.**
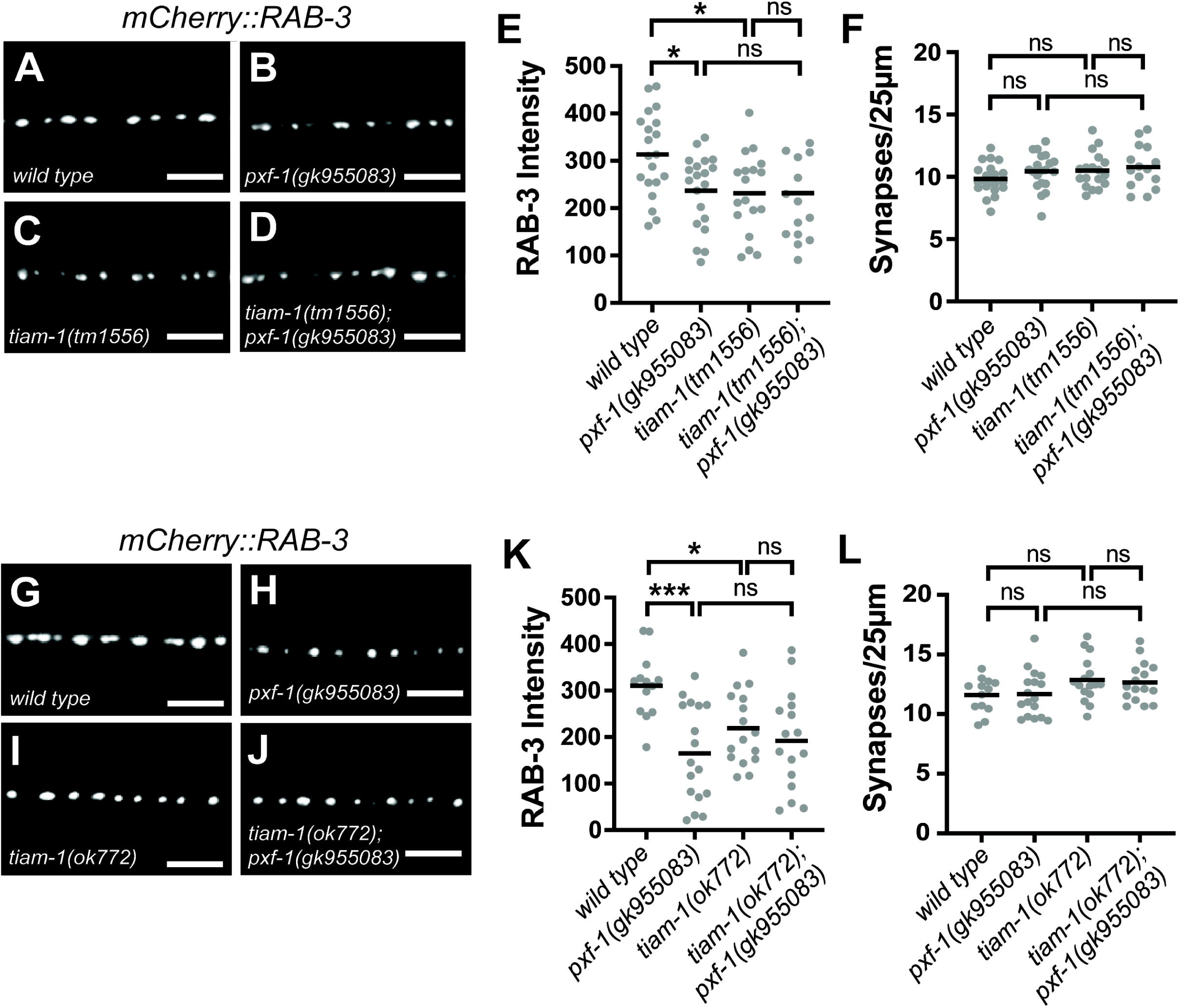
TIAM-1 functions with PXF-1 to restore synapse development. **(A-D)** Representative images of mCherry::RAB-3 labeled cholinergic synapses in the dorsal cord of **(A)** wild type, **(B)** *pxf-1(gk955083),* **(C)** *tiam-1(tm1556)*, and **(D)** *tiam-1(tm1556); pxf-1(gk955083)* animals. Scale bars are 5 μm. **(E)** Quantification of mCherry::RAB-3 grayscale fluorescence intensity. (**F)** Quantification of synaptic density. Gray dots represent individual animals, black bars indicate means. n = 15-20. *p<0.05; ns = not significant. **(G-J)** Representative images of mCherry::RAB-3 labeled cholinergic synapses in the dorsal cord of **(G)** *wild type,* **(H)** *pxf-1(gk955083),* **(I)** *tiam-1(ok772),* and **(J)** *tiam-1(ok772); pxf-1(gk955083)*. Scale bars are 5 μm. **(K)** Quantification of mCherry::RAB-3 grayscale fluorescence intensity. **(L)** Quantification of synaptic density. Gray dots represent individual animals, black bars indicate means. n = 13-17. *p<0.05; ***p<0.001; ns = not significant.

### RAP-1 interacts with TIAM-1 in cholinergic motor neurons

Based on our data, we hypothesize that PXF-1 activates RAP-1 signaling and that TIAM-1 activates RAC-2 signaling downstream of RAP-1. TIAM1 in mammals contains a Ras-binding domain, which mediates interactions between small GTPases and their downstream effectors (Lambert et al., 2002). Although *C. elegans* TIAM-1 lacks a predicted Ras-binding domain, sequence alignment among human, mouse, and *C. elegans* TIAM1/TIAM-1 proteins display ∼20% identity between *C. elegans* TIAM-1 and the Ras-binding domains from its mammalian homologs (**Figure 8A**). Therefore, we hypothesized that TIAM-1 may be a direct effector of RAP-1 in cholinergic neurons. To test this hypothesis, we used FRET-FLIM to determine if TIAM-1 and RAP-1 interact in cholinergic motor neurons. We expressed TIAM-1::mGFP with either mCherry or RAP-1::mCherry in cholinergic neurons and measured the lifetime fluorescence of mGFP (**Figure 8B**). We observed a significant decrease in TIAM-1::mGFP lifetime in the presence of RAP-1::mCherry as compared to free mCherry (**Figure 8C-E**). This decrease in TIAM-1::mGFP lifetime fluorescence is consistent with an interaction between RAP-1 and TIAM-1.

**Figure 8.**
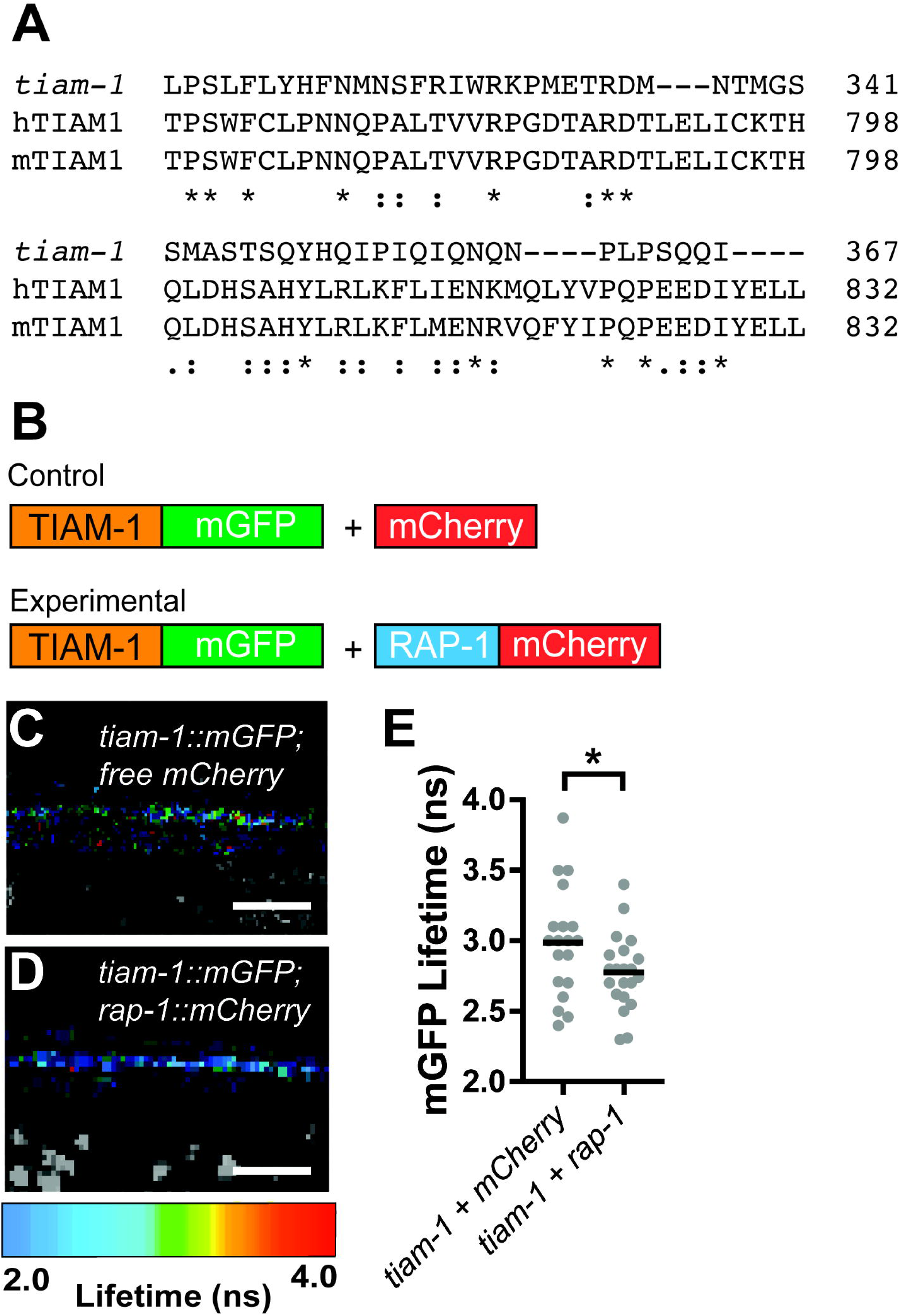
RAP-1 interacts with TIAM-1 in cholinergic motor neurons. **(A)** Sequence alignments of Ras-binding domains from human TIAM1 (hTIAM1) and mouse TIAM1 (mTIAM1) with *C. elegans* TIAM-1. Sequences were aligned using the Clustal Omega program (Sievers and Higgins, 2018; Sievers et al., 2011) through UniProt (UniProt, 2024). Degrees of amino acid conservation are indicated as an asterisk for fully conserved residues, a colon for conserved between residues with strongly similar properties, or a period for conserved between residues with weakly similar properties. **(B)** Schematic of plasmids expressing TIAM-1::mGFP and RAP-1::mCherry in cholinergic motor neurons used for FRET-FLIM experiments. **(C & D)** FastFLIM images of mGFP in the dorsal cord of animals expressing each combination of plasmids in cholinergic neurons. The lifetime for mGFP has been pseudo-colored to represent fluorescence lifetime values in nanoseconds. Scale bar is 5 μm. **(E)** Mean lifetime fluorescence of TIAM-1::mGFP *in vivo* with control versus experimental plasmids. Data was pooled from multiple transgenes: TIAM-1::mGFP + mCherry (*bluEx148*, *bluEx149,* and *bluEx150*) and TIAM-1::mGFP + RAP-1::mCherry (*bluEx152, bluEx153,* and *bluEx154*). Gray dots represent individual animals, black bars indicate means. n = 19-21. *p<0.05.

## Discussion

Synapse development relies on precisely coordinated signaling events to establish and maintain functional connections. Small G proteins can localize to specific regions of cells and oscillate between active and inactive states. Therefore, modulation of their localization or duration of activity represents one way to coordinate the growth and development of tissues. In the nervous system, Rap proteins influence synaptic connectivity and function by modulating the density of dendritic spines, abundance of neurotransmitter receptors, or numbers of synaptic terminals (Beaudoin et al., 2012; Chen et al., 2018a; Heo et al., 2017; Ryu et al., 2008; Zhu et al., 2002). Here, we show that *C. elegans* RAP-1, a target of PXF-1, promotes the development of presynaptic terminals at the neuromuscular junction. Our data indicate that PXF-1 induction of RAP-1 signaling leads to the activation of RAC-2, which modulates the presynaptic actin cytoskeleton. We propose that activated RAP-1 interacts with TIAM-1 to connect RAP-1 signaling to RAC-2.

In mammalian cells, a similar interaction between Rap and Rac GTPases promotes cytoskeletal regulation (Taira et al., 2004). Multiple studies outside the nervous system support a direct interaction between Rap1 and two Rac GEFs, Tiam1 or VAV2, to modulate actin filaments in a Rac-dependent manner (Arthur et al., 2004; Birukova et al., 2013; Birukova et al., 2008; Gérard et al., 2007). Similar pathways also underlie cortical development. For example, Rap1 acts upstream or parallel to Rac1 to promote neuronal migration (Jossin and Cooper, 2011). This study suggested that Vav2 activation is the most likely Rac GEF to connect Rap1 to Rac GTPase signaling during cortical neuron migration. Our study suggests that RAP-1 acts through TIAM-1 and Rac GTPase to promote the accumulation of synaptic vesicles at presynaptic terminals. However, we have not investigated which pathways may be involved in the increased synaptogenesis observed in *rap-1* mutants. Since the small GTPases and regulatory proteins investigated here do not display a synaptogenesis phenotype, we would hypothesize that the role of RAP-1 during synaptogenesis may proceed through a different Rac GEF or set of effector proteins.

### Divergent roles of Rap GTPases at the synapse

Throughout evolution, most species contain at least two Rap paralogs, Rap1 and Rap2. While both paralogs facilitate similar biological processes, they appear to operate through different signaling mechanisms or may even oppose one another. For example, Rap1 in mammals promotes dendritic spine elongation in cortical neurons, but Rap2 activation causes the formation of shorter spines in the forebrain of mice (Ryu et al., 2008; Xie et al., 2005). In hippocampal neurons, activation of Rap2, but not Rap1, reduces the complexity of axons and dendritic arbors; however, inhibition of Rap1 signaling resulted in shorter dendrites (Fu et al., 2007). These findings suggest that differential activation of Rap1 and Rap2 is important for dendritic development.

In *C. elegans* cholinergic motor neurons, *rap-2* prevents the overlap of adjacent presynaptic regions but does not alter overall synapse number (Chen et al., 2018a). Here we show that *rap-1* promotes synaptic vesicle marker intensity and reduces the density of cholinergic synapses, which are not altered by *rap-2* (**Figure 2**). The divergent effects of these Rap paralogs at the synapse in some cases can be attributed to different signaling pathways. At postsynaptic sites for example, Rap1 acts through AF-6 to influence dendritic spine neck length, but Rap2 opposes ERK signaling to reduce spine length (Ryu et al., 2008; Xie et al., 2005). At presynaptic terminals, *C. elegans* RAP-2 signals through TNIK to modulate the actin cytoskeleton (Chen et al., 2018a). Identifying the mechanisms that differentially modulate Rap paralogs in presynaptic and postsynaptic compartments should reveal how these distinct signaling pathways converge on synapse development and function.

### Regulation of Rac activity

Activation of RAP-1 was required to rescue *pxf-1* mutant deficits in synapse development; however, simply overexpressing RAC-2 was sufficient to restore synapse development in *pxf-1* mutants (**Figure 6**). This suggests that exogenous RAC-2 can be activated in *pxf-1* mutants to overcome the lower level of endogenous RAC-2 activity in *pxf-1* mutants. Since *tiam-1* and *pxf-1* mutants cause similar decreases in RAC-2 activity (**Figure 5**), we would infer that *pxf-1* mutants do not display a compensatory increase in TIAM-1 activity, which could activate exogenous RAC-2 to restore RAC-2 activity levels to wild type. Therefore, we hypothesize that while some exogenous RAC-2 is activated by TIAM-1, some RAC-2 may also act as a decoy for Rac GAPs, allowing GTP-bound RAC-2 to reach wild type levels in *pxf-1* mutants. While we have not tested this idea, similar effects were observed in mammalian cells using a Rac GAP dominant negative mutation. For example, overexpression of a dominant negative Rac GAP, Bcr, in mammalian neurons increases Rac activity to a similar level as Tiam1 overexpression (Um et al., 2014). There are no predicted homologs for Bcr or its close relative, Abr, in *C. elegans.* However, identification of the Rac GAP that acts with TIAM-1 to balance RAC-2 activity will allow for further investigation of the signaling between RAP-1 and RAC-2 during synapse development. It also will enable us to more precisely measure how GEFs and GAPs contribute to G protein activation levels *in vivo*. Additional studies will be required to determine how RAP-1 regulates TIAM-1 function.

### Regulation of Rap GEF activity

GAPs and GEFs coordinate where and when G proteins are active. While protein-protein interactions oftentimes determine where GAPs and GEFs influence GTPase activity, it’s not always clear if and how the activity of GAPs or GEFs is regulated. Some Ras and Rap GEFs are activated by second messengers, like calcium, diacylglycerol, or cAMP (de Rooij et al., 2000; de Rooij et al., 1998; Ebinu et al., 1998; Kawasaki et al., 1998). Exchange-protein activated by cAMP (Epac) and Ras-associating (RA)-GEF/PDZ-GEF proteins constitute two very similar classes of Rap GEFs. Both contain cyclic nucleotide-binding domains, yet only Epacs are known to be activated by cAMP binding (de Rooij et al., 2000; de Rooij et al., 1998; Kawasaki et al., 1998). RA-GEF/PDZ-GEFs, like *pxf-1*, RAPGEF2, or RAPGEF6, do not appear to be modulated by cAMP or cGMP (Liao et al., 1999). Additional studies suggest that binding of Ras or Rap to these RA-GEF/PDZ-GEFs acts as positive feedback loop to amplify the activity of these G proteins (Hu et al., 1999; Jin et al., 2001; Rebhun et al., 2000). Moreover, post-translational modification of Rap GEFs has been shown to modulate Rap GEF activity during neuronal migration (Ye et al., 2014); however, it is not known whether similar events occur during synapse development. Further studies will be required to determine if PXF-1 and its homologs are modulated by post-translational modifications, the binding of cyclic nucleotide monophosphates, or additional G proteins during synapse development.

## Materials and methods

### DNA Cloning and Plasmid Construction

To express rap-1 in cholinergic neurons, *rap-1(WT)* cDNA was cloned by PCR with Q5 polymerase (NEB) using oSC188 and oSC189. We cloned *rap-1(G12V)* cDNA from pREA6 [*Prgef-1::rap-1 (G12V) cDNA*] using oSC188 and oSC189. Using the Gibson assembly method, we inserted the *rap-1(WT)* PCR product into pBD23 [*pCR8-mGFP*] digested with KpnI (NEB) to create pBD25 [*pCR8 rap-1(WT)::mGFP*] and the *rap-1(G12V)* PCR product into pBD28 [*pCR8 tiam-1::mGFP*] digested with KpnI to create pSJC316 [*pCR8 rap-1(G12V)::mGFP*]. Using LR clonase II (Invitrogen), we inserted *rap-1(WT)::mGFP* or *rap-1(G12V)::mGFP* into pCZGY1091 [*Punc-17b-dest*] to create pBD31 [*Punc-17b::rap-1(WT)::mGFP*] and pSJC319 [*Punc-17b::rap-1(G12V)::mGFP*].

To express rac-2 in cholinergic neurons, we cloned *rac-2(WT)* or *rac-2(G12V)* cDNA from pREA2 [*Prgef-1::rac-2(WT) cDNA*] or pREA3 [*Prgef-1::rac-2(G12V) cDNA*], respectively, by PCR with Q5 polymerase using oSC304 and oSC305. The *rac-2(WT)* or *rac-2(G12V)* PCR products were inserted into pBD28 [*pCR8 tiam-1::mGFP*] digested with KpnI to create pSJC317 [*pCR8 rac-2(WT)::mGFP*] and pSJC318 [*pCR8 rac-2(G12V)::mGFP*]. Using LR clonase II, we inserted the *rac-2(WT)::mGFP* or *rac-2(G12V)::mGFP* into pCZGY1091 [*Punc-17b-dest*] to create pSJC320 [*Punc-17b::rac-2(WT)::mGFP*] and pSJC321 [*Punc-17b::rac-2(G12V)::mGFP*].

For cholinergic expression of the RAC-2 activity biosensor, we created pREA12 [*pCR8 mCherry::RBD::T2A::rac-2::mGFP]* using Gibson assembly between pSJC213, a modified pCR8 Gateway entry vector, digested with NdeI (NEB) and 2 gBlock (Integrated DNA Technologies) encoding *mCherry::RBD::rac-2 N-terminus*, which includes the Rac binding domain (RBD) from *pak-1* and a T2A ribosomal skip sequence, and *rac-2 C-terminus::mGFP*. pREA16 [*Punc-17b::mCherry::RBD::T2A::rac-2::mGFP*] was generated using an LR reaction between pREA12 and pCZGY1091.

To measure protein-protein interactions between RAP-1::mCherry and TIAM-1::mGFP, we created pBD28 [*pCR8 tiam-1::mGFP*] using Gibson assembly between 2 gBlocks encoding the N-terminus and C-terminus of *tiam-1* cDNA and pBD23 digested with KpnI. pBD24 [*pCR8 rap-1::mCherry*] was created by Gibson assembly between pBD22 [pCR8 mCherry] digested with KpnI and *rap-1* cDNA amplified by PCR as described above. We used LR clonase II to insert *tiam-1::mGFP* or *rap-1::mCherry* into pCZGY1091 to create pBD34 [*Punc-17b::tiam-1::mGFP*] and pBD30 [*Punc-17b::rap-1::mCherry*]. All entry vectors sequences were confirmed by Sanger sequencing (Eurofins). Plasmid design and sequencing results were analyzed using ApE (Davis and Jorgensen, 2022).

### C. elegans strains, maintenance, and transgenesis

All studies were performed using *C. elegans* hermaphrodites maintained on standard nematode growth medium plates (NGM) at 20°C and seeded with OP50 (*Caenorhabditis* Genetics Center; WormbaseID: OP50) as previously described (Brenner, 1974). All transgenic strains were generated in the lab as using the previously described microinjection technique (Mello et al., 1991) using *Pmyo-2::mCherry* (2ng/μl) or *Punc-122::RFP* (60ng/μl) co-injection markers. Details about the strains generated for this study are listed in Table 1. Genetic mutants were identified by genotyping with the following primer sets: oSC48 and oSC49 for *pxf-1(gk955083)*, oSC236 and oSC237 for *rap-1(pk2082)*, oRL21, oRL22, and oRL23 for *rap-2(gk11)*, oRL4, oRL5, and oRL6 for *rac-2(ok326)*, oBM65, oBM66, and oBM63 for *tiam-1(tm1556)*, oBM62, oBM63, and oBM64 for *tiam-1(ok772)*. Primer sequences are listed in Table 2.

**Table 1:**
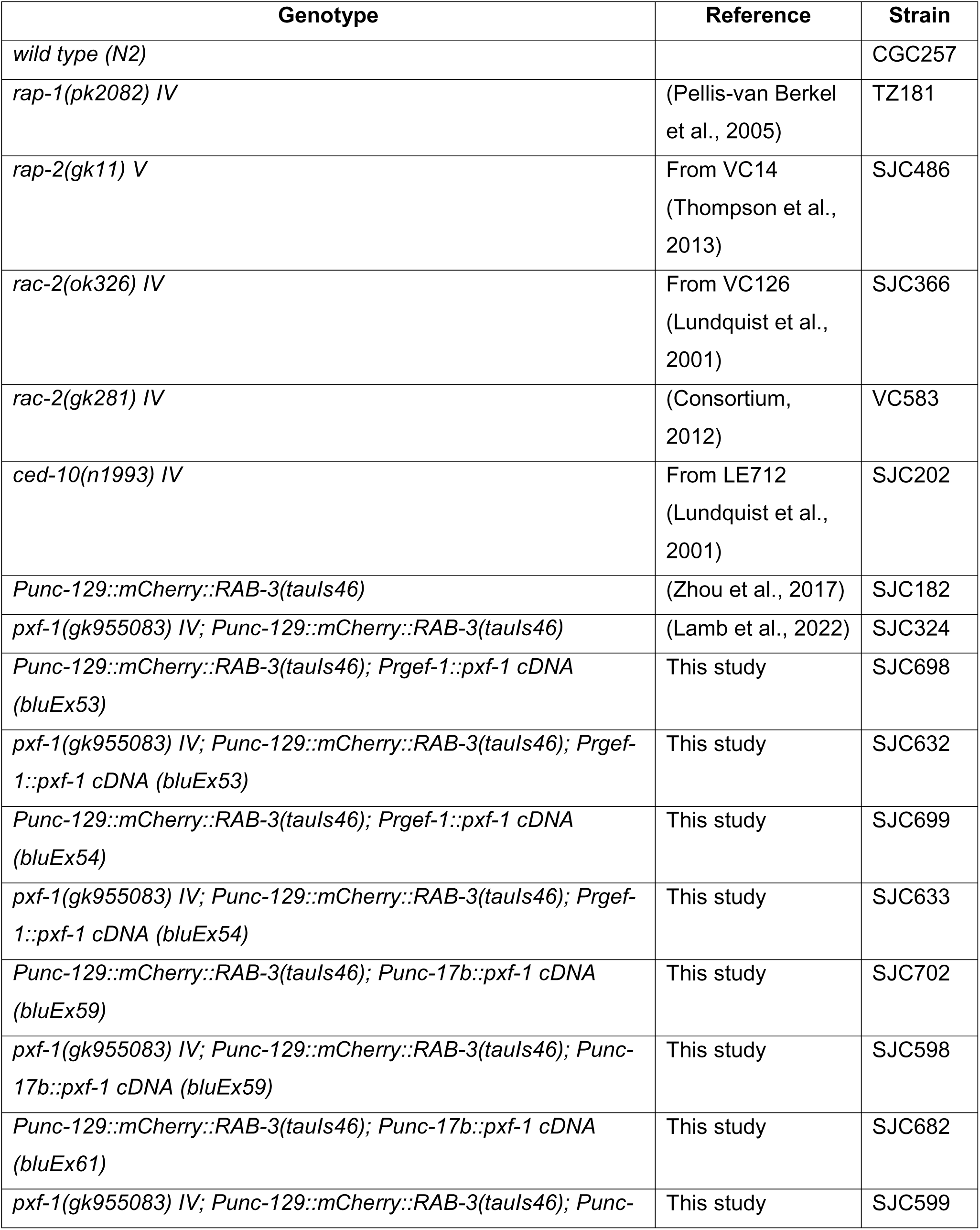
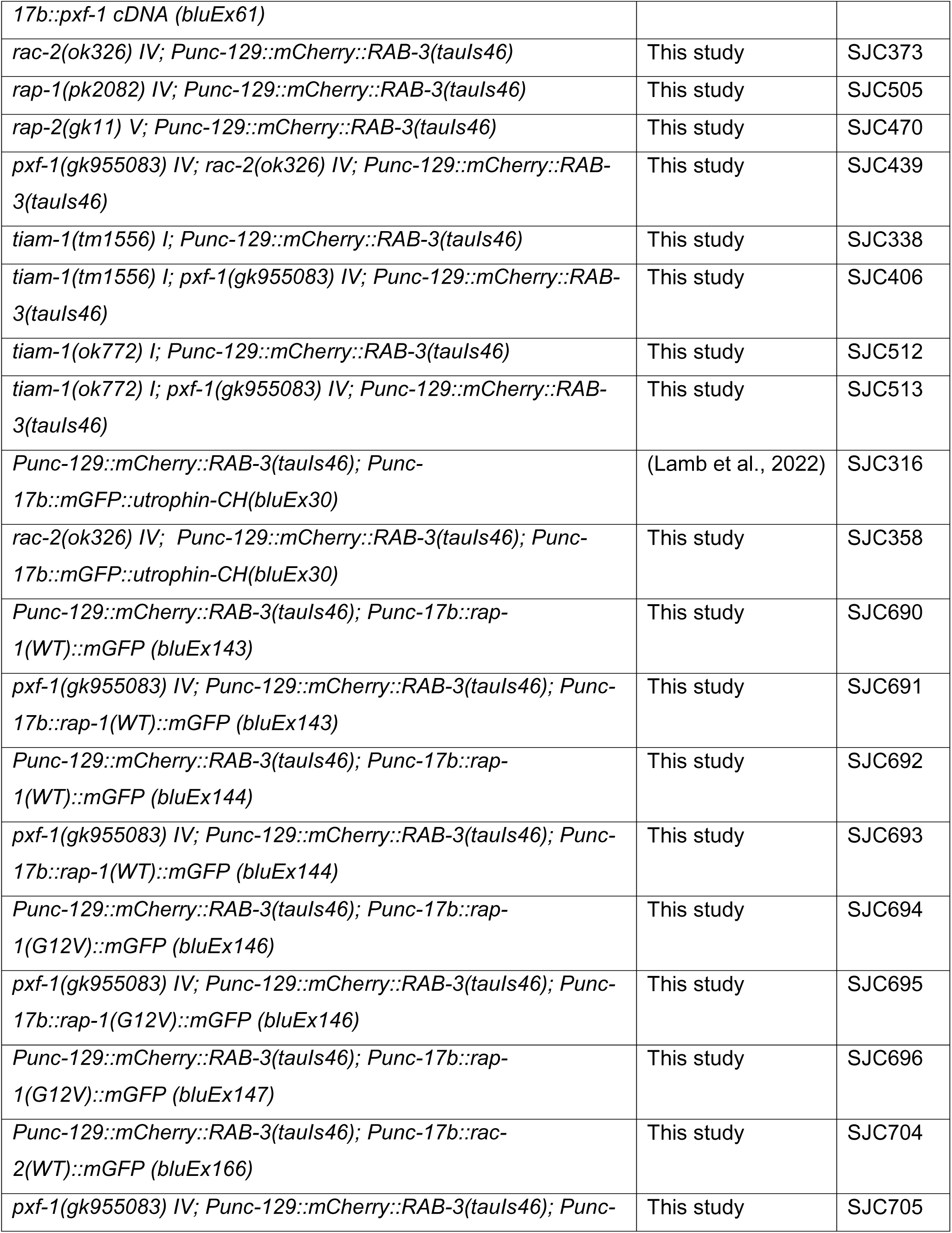
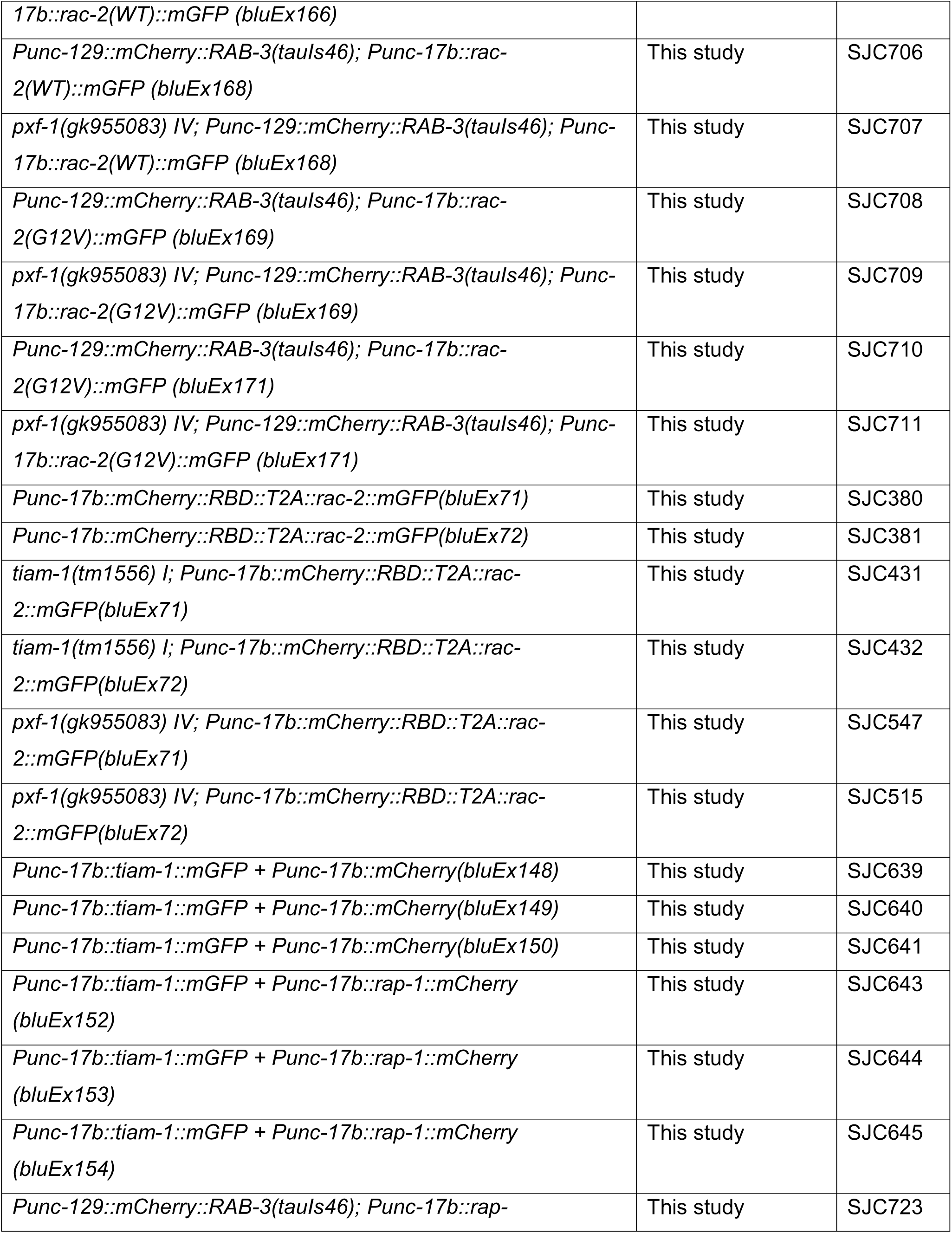
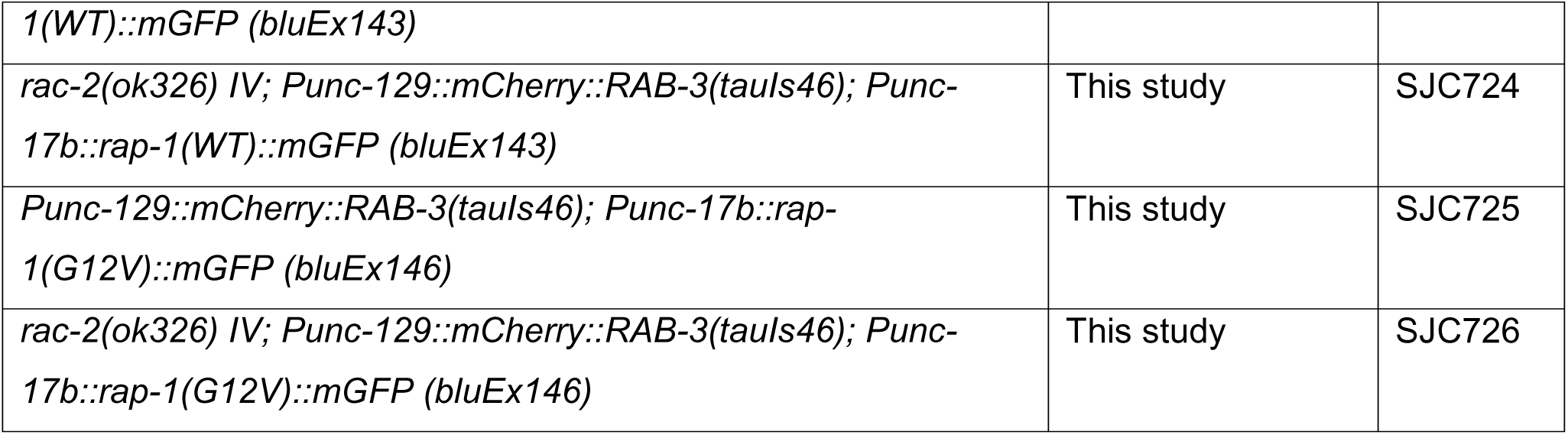
*C. elegans* strains used in this study.

**Table 2:**
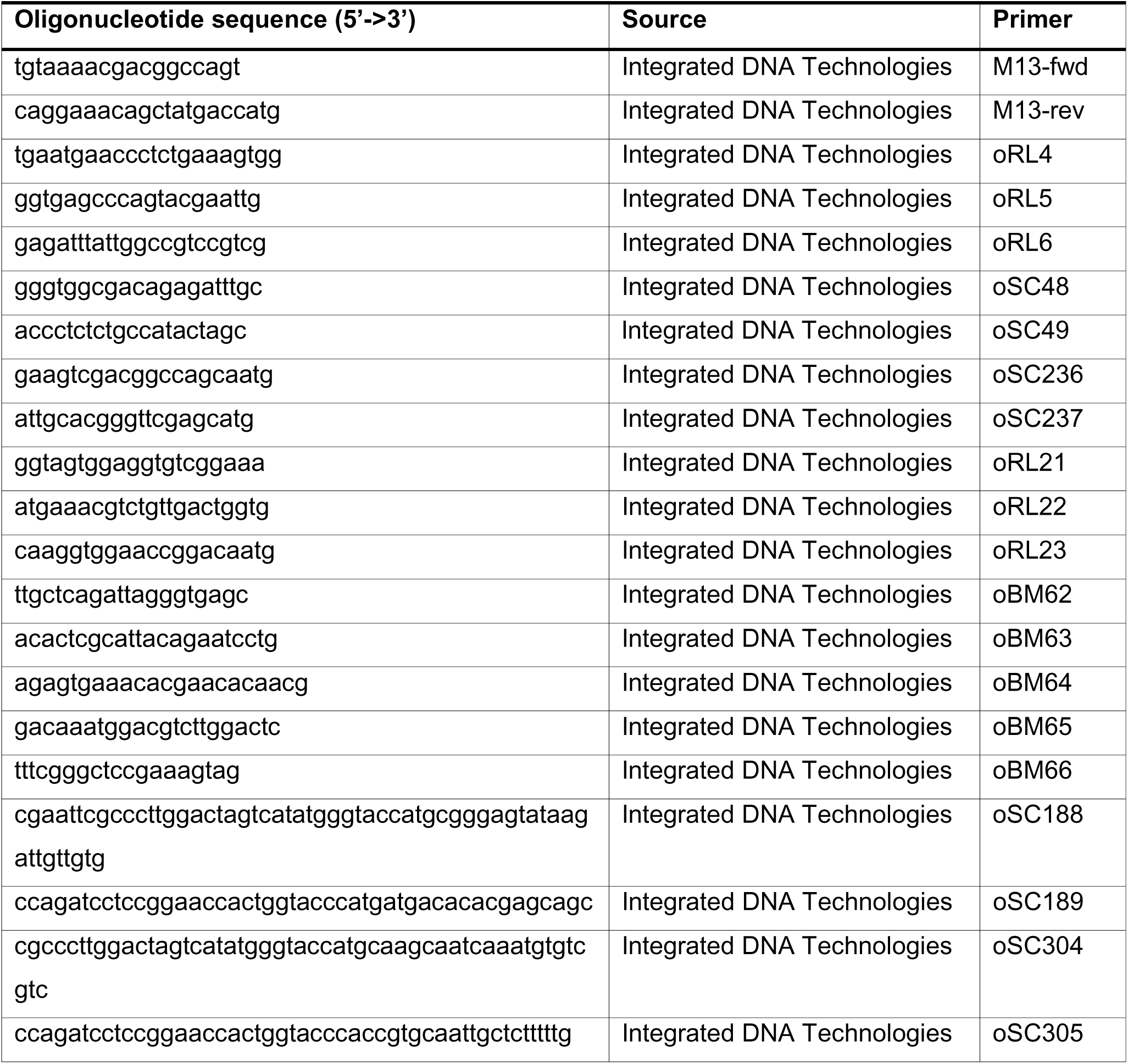
Oligonucleotides used in this study.

### Aldicarb Behavioral Assay

Day one adults were placed on NGM plates containing 1mM aldicarb (Sigma-Aldrich) and scored for paralysis at 15-minute intervals during two hours of observation. Plates were prepared and poured in batches and kept at 4°C prior to use. Paralysis was defined as no response after three touches. Plates were placed on the bench 30min before the experiments, and all assays were performed at room temperature (20-21°C). Individuals performing the assays were blinded to the genotypes.

### Fluorescence Microscopy

To visualize changes to synaptic vesicle pools, mutant animals were crossed with mCherry::RAB-3(*tauIs46)* synaptic vesicle marker. To visualize filamentous actin, mutant animals were crossed with mGFP fused to the calponin homology domain of utrophin (*bluEx30*) (Lamb et al., 2022). Animals were immobilized in M9 buffer on 10% agarose pads. Images were taken with a DS-Qi2 camera at 60X magnification with a 1.20 NA water immersion lens using Ti-2E widefield fluorescent microscope (Nikon) along the posterior dorsal cord in L4 animals and analyzed using the FIJI distribution of NIH ImageJ (Schindelin et al., 2012) as previously described (Lamb et al., 2022).

### Whole Mount Immunohistochemistry

To visualize endogenous expression of UNC-17, adult animals were fixed as previously described (Cherra and Jin, 2016, Lamb et al. 2022). Samples were blocked with 5% BSA for 1 hr and incubated overnight with an UNC-17 antibody (1:500 dilution, Mab1403 Developmental Studies Hybridoma Bank, AB_2315531). Samples were washed with 2% BSA in PBST and incubated for 1 hr with Alexa Fluor 594 donkey anti-mouse secondary antibody (Thermo Scientific, AB_141633). After washing, samples were mounted using Vectashield mounting medium (VWR), covered with a coverslip, and imaged using the Ti2-E microscope (Nikon) at 60x magnification with 1.20NA water immersion lens (Nikon).

### Fluorescence Lifetime Imaging Microscopy

To visualize *in vivo* changes to RAC-2 activity or interactions between RAP-1 and TIAM-1, L4 animals containing FRET-FLIM biosensors expressed in cholinergic neurons were immobilized in M9 buffer on 10% agarose pads. FRET-FLIM biosensors were constructed using the mGFP-mCherry donor acceptor pairing under the *unc-17b* promoter. Cholinergic synapses were located along the dorsal cord using the FITC laser on a Nikon A1 Confocal microscope. Images were acquired for 1 min at 20°C using a 60x 1.20NA water immersion lens (Nikon) using a PMA 40 Hybrid Photomultiplier Detector (Picoquant) and a SuperK Extreme white light laser (NKT Phototonics) equipped with a 488 nm filter. Laser was pulsing at 38.9 MHz. Output signal was measured using a Picoharp 300 Time-Correlated Single Photon Counting system at 8.0 ps resolution (Picoquant). All frames were binned together. Mean fluorescence lifetimes of mGFP molecules were calculated using PicoQuant SymPho Time 64 software.

### Data quantification and statistical analysis

All statistical analyses were performed using Prism 9 (GraphPad). For comparisons between two groups, we used a *t*-test when distributed normally and a Mann-Whitney test for non-parametric data. For comparisons between multiple groups with normally distributed data, a one-way ANOVA was performed followed by Sidak’s multiple comparisons test. For data that was not normally distributed, a Kruskal-Wallis test was used followed by Dunn’s multiple comparisons test. We used a Mixed-effects analysis followed by Dunnett’s multiple comparisons for drug-induced paralysis time courses. A corrected *p*-value less than 0.05 was considered significant. Additional statistical information for individual figures may be found within the legends.

## Acknowledgments

We would like to thank Avanti Sawardekar and Bithika Dhar for the creation of reagents and initial validation of cDNA expression. FRET-FLIM image acquisition and data analysis were performed using resources at the Light Microscopy Core at the University of Kentucky. Some mutant strains were provided by the *Caenorhabditis* Genetics Center, which is supported by NIH (OD010440).

